## Supplementary material for "Intrinsically disordered protein biosensor tracks the physical-chemical effects of osmotic stress on cells": Figure S

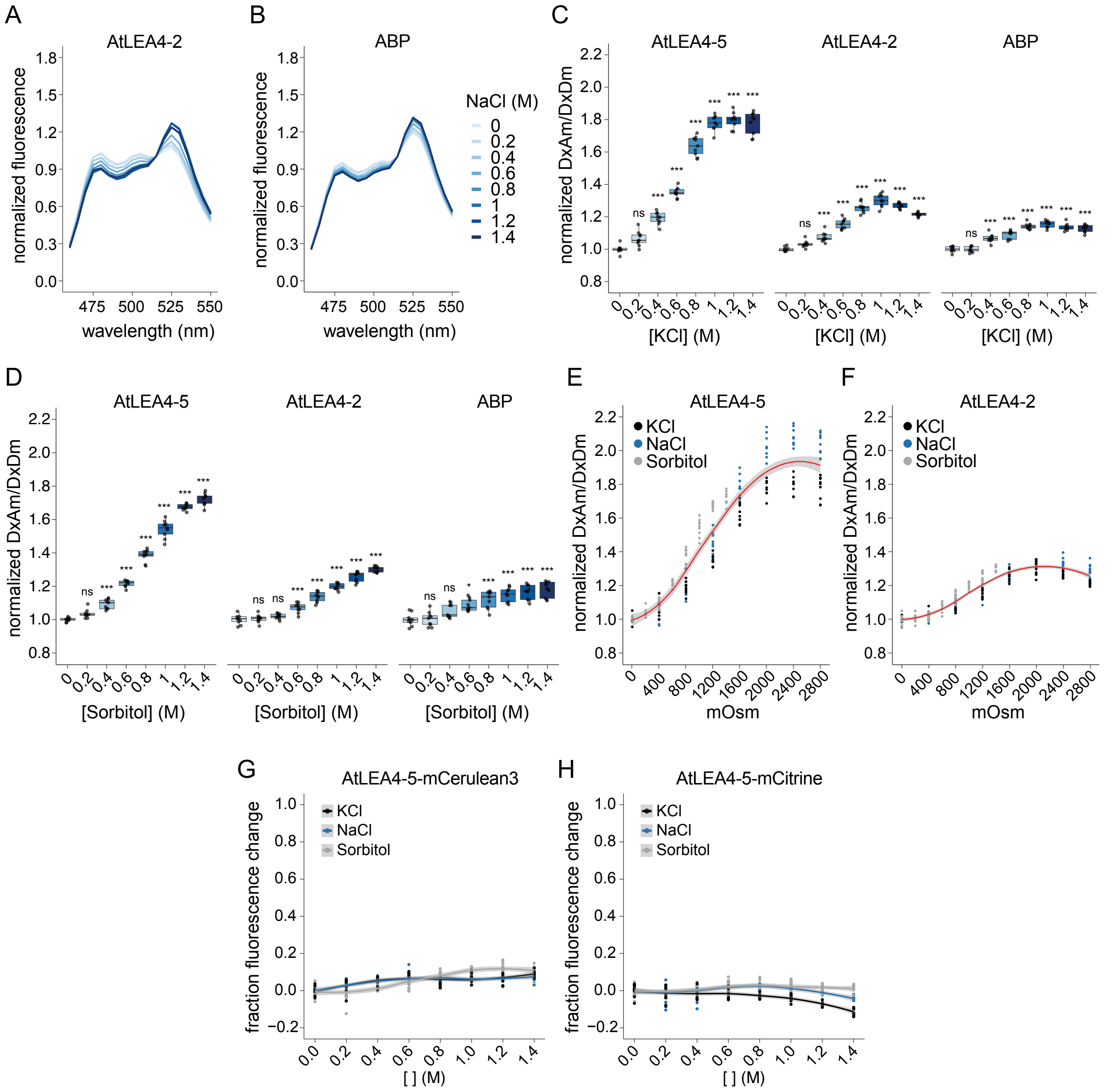


**Figure S1.** (A) Fluorescence emission spectra of NaCl-treated live yeast cells expressing the biosensor construct using AtLEA4-2 as the sensory domain. Fluorescence values were normalized to the value at 515 nm. (B) Fluorescence emission spectra of NaCl-treated live yeast cells expressing the biosensor construct using ABP as the sensory domain. Fluorescence values were normalized to the value at 515 nm. (C) Normalized FRET ratio (DxAm/DxDm) of live yeast cells treated with different concentrations of KCl. Cells are expressing the biosensor construct using either AtLEA4-5, AtLEA4-2, or ABP as the sensory domain. Two-way ANOVA. *p < 0.05, **p < 0.01, ***p < 0.001. (D) Normalized FRET ratio (DxAm/DxDm) of live yeast cells treated with different concentrations of sorbitol. Cells are expressing the biosensor construct using either AtLEA4-5, AtLEA4-2, or ABP as the sensory domain. Two-way ANOVA. *p < 0.05, **p < 0.01, ***p < 0.001. (E) Normalized FRET ratio (DxAm/DxDm) of live yeast cells expressing the biosensor construct using AtLEA4-5 as the sensory domain. Cells were hyperosmotically shocked to the indicated osmolarity with either NaCl (blue), KCl (black), or sorbitol (gray). (F) Normalized FRET ratio (DxAm/DxDm) of live yeast cells expressing the biosensor construct using AtLEA4-2 as the sensory domain. Cells were hyperosmotically shocked to the indicated osmolarity with either NaCl (blue), KCl (black), or sorbitol (gray). (G) Fraction fluorescence change relative to the standard condition of live yeast cells expressing AtLEA4-5-mCerulean3 (donor-only construct) when cells were exposed to different concentrations of NaCl (blue), KCl (black), or sorbitol (gray). (H) Fraction fluorescence change relative to the standard condition of live yeast cells expressing AtLEA4-5-mCitrine (acceptor-only construct) when cells were exposed to different concentrations of NaCl (blue), KCl (black), or sorbitol (gray).

A

| Construct | Donor | Acceptor |
|---|---|---|
| FP1 | t7.eCFP.t9 | Aphrodite.t9 |
| FP2 | t7.TFP.t9 | Aphrodite.t9 |
| FP3 | mTFP.t9 | Aphrodite.t9 |
| FP4 | Cerulean | Aphrodite.t9 |
| FP5 | Cerulean | Citrine |
| FP6 | edCerulean | edCitrine |
| FP7 | t7.eCFP.t9 | edAphrodite.t9 |
| FP8 | Citrine | mCerulean |
| FP9 | edCerulean | edAphrodite.t9 |
| FP10 | mCerulean3 | mCitrine |

B

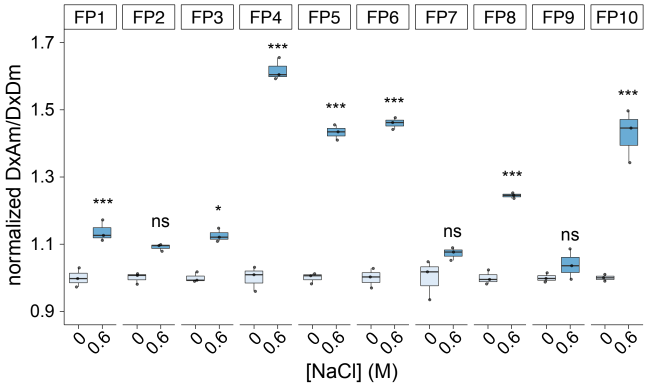


**Figure S2.** (A) Different FRET pairs used in this study. FP: FRET pair. t7: 7 amino acid terminal truncation. t9: 9 amino acid terminal truncation. m: monomeric variant. ed: enhanced dimerization variant. (B) Normalized FRET change of live yeast cells expressing AtLEA4-5 fused to the different FRET pairs in (A), subjected to 0 M or 0.6 M NaCl. Two-way ANOVA. *p < 0.05, **p < 0.01, ***p < 0.001.

A

**AtLEA4-5**

```
MQSMKETASN IAASAKSGMD KTKATLEEKA EKMKTRDPVQ KQMATQVKED
KINQAEMQKR ETRQHNAAMK EAAGAGTGLG LGTATHSTTG QVGHGTGTHQ
MSALPGHGTG QLTDRVVEGT AVTDPIGRNT GTGRTTAHNT HVGGGGATGY
GTGGGYTG
```

**Scramble-1**

```
VHDGETPVGP RTEQAMTTQE QEAKNTGGLA NIGAQTDTKK QTAQASHQTA
TEKTKAKGAT GRRTGGYGNN HGMVKMGVIM GSKGKEMGLR TDTMTTGHSS
GIGKQHLMAL RHAATMGTRA SSGVKGGETV THEPAAAGQG KGTNYLATTG
EQADVGTD
```

**Scramble-2**

```
GPSIKKTQTL KHPGNALEMT VGMAEGGNTV ARTHDSSGDS HQTAETKDHG
GTGAMYKLHK HNGKTQQVDT GMATAGAVPY AKEKTSTGSN ATAEHTTGKM
TGTAQIMETG RLAERVGMTI DARTGGLNGG GTARMQRKQT QGAEKAGVGQ
QAGVTGET
```

**Scramble-3**

```
GDTNGSTTAM RAQTDEKMAT RQKEAGNDRY TSGHTAGVTA KTLGLPDARS
KSGPKKSTET GEHQGTMEQY TTHGEGHGKA AQVIAATGLK TNTKQANGIK
KTLQRQMQTA VGAMNHEGEA AVALVGGHGE HAGTVVGGTQ GTGMTMSMGI
PTDKTGGR
```

**Scramble-4**

```
QGGMGKAHKG EQKGTVTTNI AVGAVTAGHQ TEGLEQGGHE ADTVAAMAKS
LQPKRTTTGT GNTRTLELSG HTAATNSMPK GTHEMGEMTA QGRIGTMVKL
GGYQSGTHGE DAQAKKAGDS GSTKRDNAGA RQIKAETKVG VMGTHTTTQD
RANYGPMT
```

**Scramble-5**

```
SHPAGMKATM GHQHEYQEQT SAMGKGTLEI EDTGTAAGEQ RTHDKASTGA
TTTGDQETMG GMQAKVKMHL LTAGLQAVGE AGATTMREHY GTHPTAGQNV
RPKGTTQGSK ADRILDTRMN SVKKAGGEIN GTNSGGGGTK VGAKTTNTGA
KGVRQATV
```

B

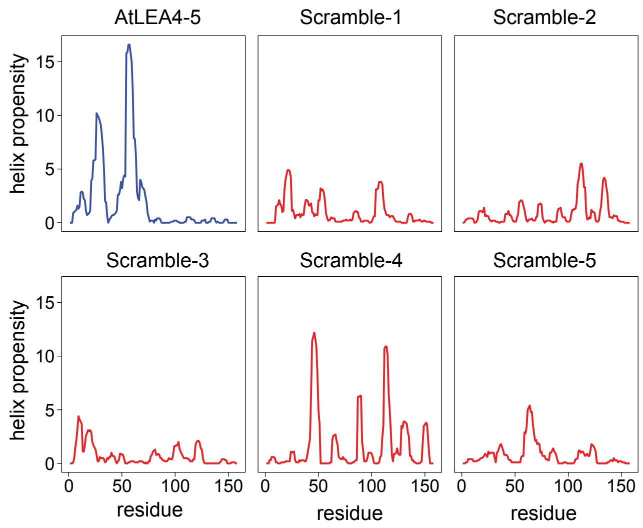


**Figure S3.** (A) Amino acid sequence of AtLEA4-5 and the five different scrambled versions of AtLEA4-5 used in this study. (B) Agadir $\alpha$-helix propensity (%) prediction of AtLEA4-5 (blue) and the five different scrambles of AtLEA4-5 (red).

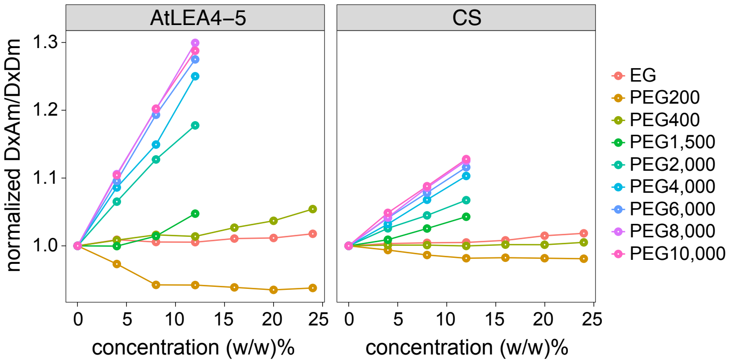


**Figure S4.** (A) Normalized FRET ratio (DxAm/DxDm) of purified recombinant full-length AtLEA4-5 or CS biosensor constructs at different concentrations (% w/w) of ethylene glycol (EG) or the indicated molecular weight polyethylene glycol (PEG).

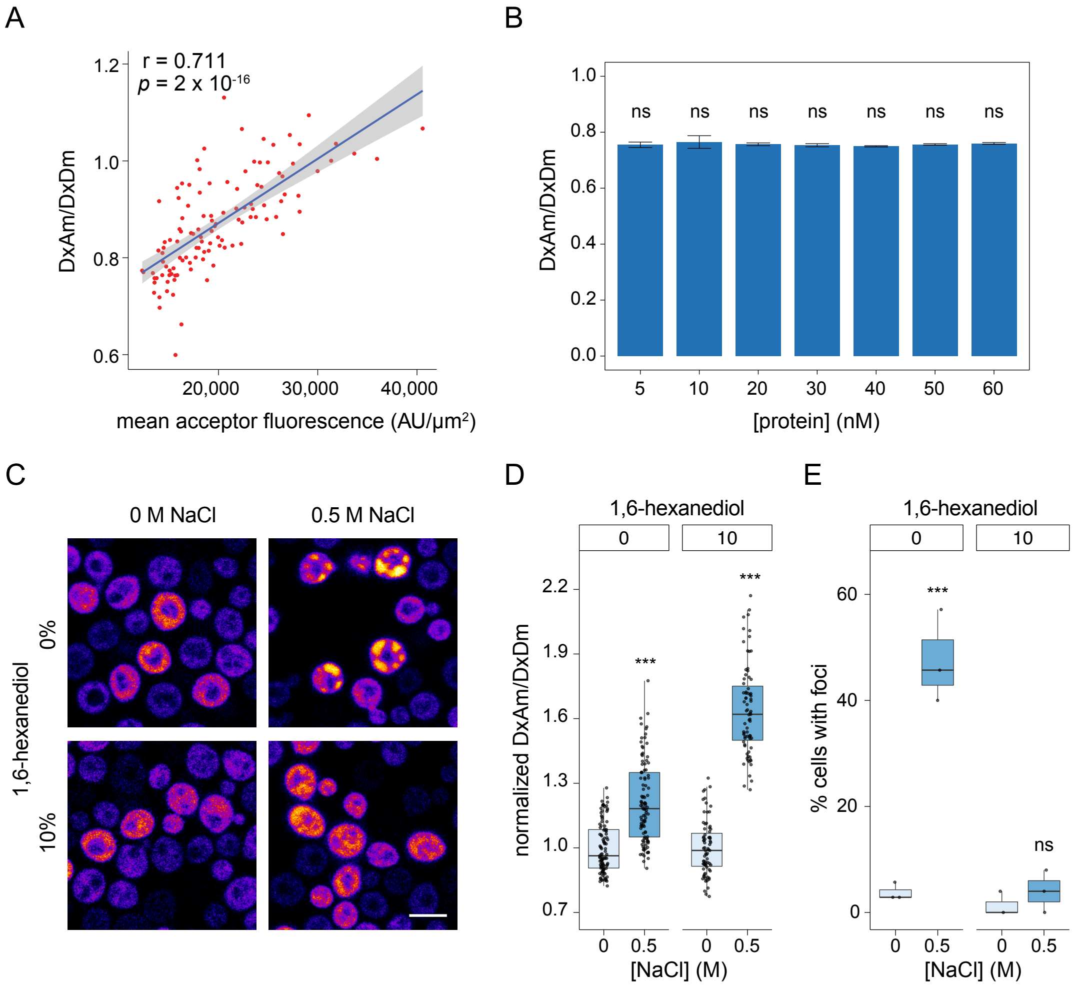


**Figure S5.** (A) Pearson's correlation of FRET ratio (DxAm/DxDm) and mean acceptor fluorescence intensity (AU/$\mu m^2$) of individual live yeast cells. r = 0.711, p-value = $2 \times 10^{-16}$. (B) FRET ratio (DxAm/DxDm) of purified recombinant full-length SED1 at different concentrations. Mean ± SEM. One-way ANOVA. *p < 0.05, **p < 0.01, ***p < 0.001. (C) Acceptor emission channel with direct acceptor excitation (AxAm) of SED1-expressing live yeast cells under 0 M NaCl and 0.5 M NaCl treatments, in the presence or absence of 10% (w/v) 1,6-hexanediol. Scale bar = 5 $\mu$m. (D) Normalized FRET ratio (DxAm/DxDm) of SED1-expressing live yeast cells under 0 M NaCl and 0.5 M NaCl treatments, in the presence or absence of 10% (w/v) 1,6-hexanediol. Two-way ANOVA. *p < 0.05, **p < 0.01, ***p < 0.001. (E) Percentage of cells with visible foci in SED1-expressing live yeast cells under 0 M NaCl and 0.5 M NaCl treatments, in the presence or absence of 10% (w/v) 1,6-hexanediol. Two-way ANOVA. *p < 0.05, **p < 0.01, ***p < 0.001.

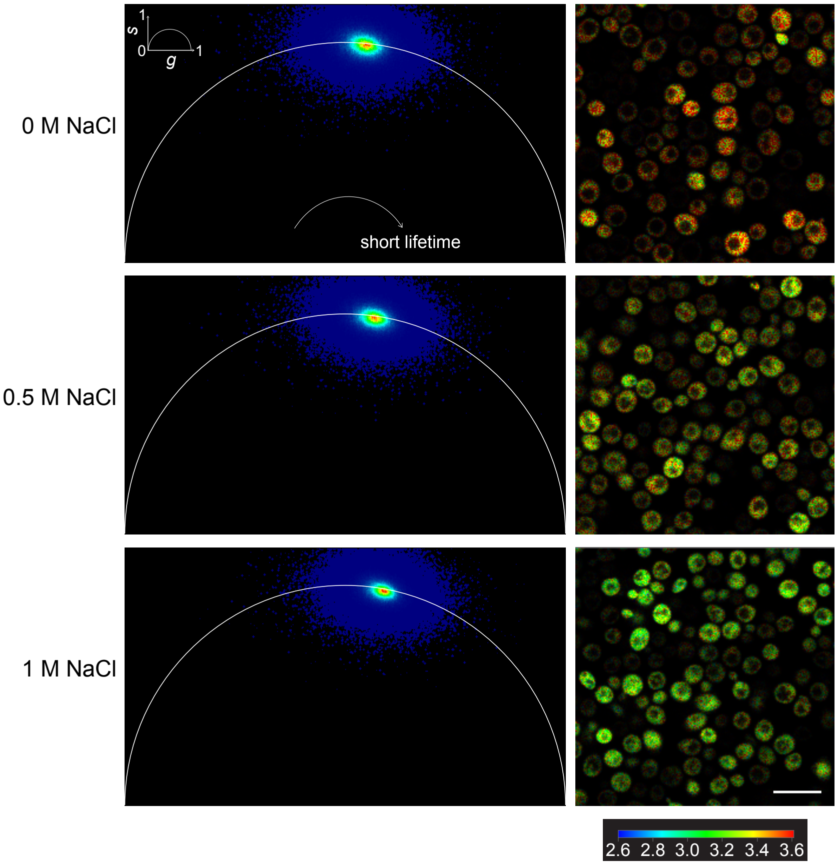


**Figure S6.** Phasor plots (left) and donor fluorescence lifetime images (right) of live yeast cells expressing AtLEA4-5 fused to mCerulean3 (donor-only control) under 0 M, 0.5 M, and 1 M NaCl. Signals shifted to the left side of the phasor plot represent longer fluorescence lifetimes, whereas those shifted to the right siderepresent shorter fluorescence lifetimes. Scale bar = 10 $\mu$m. Calibration bar represents the donor fluorescence lifetime in nanoseconds (ns).

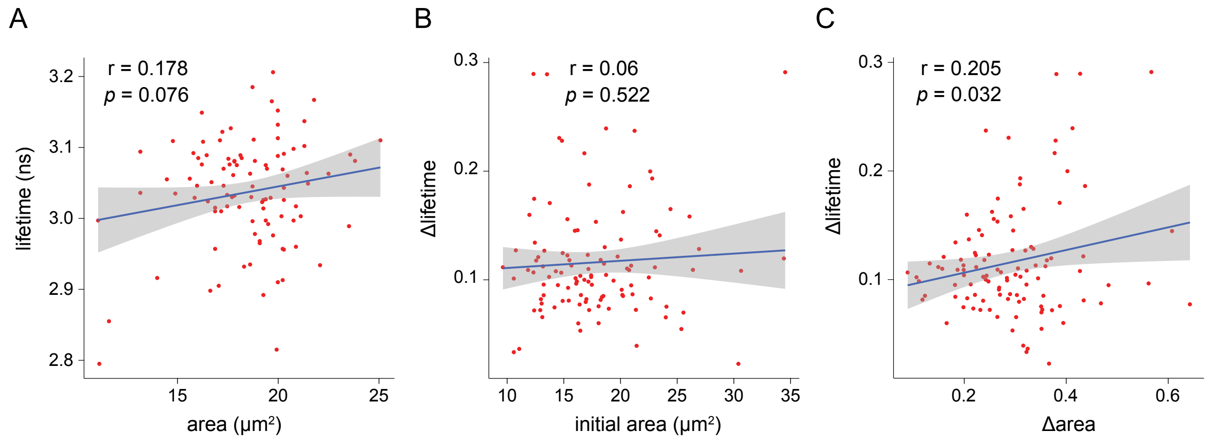


**Figure S7.** (A) Pearson's correlation of donor fluorescence lifetime (ns) and yeast area (proxy of cell volume) under standard conditions (0 M NaCl). r = 0.178, p-value = 0.076. (B) Pearson's correlation of the change in donor fluorescence lifetime (Δlifetime) and the area (proxy of cell volume) of single yeast cells before treatment with 1 M NaCl. r = 0.06, p-value = 0.522. (C) Pearson's correlation of the change in donor fluorescence lifetime (Δlifetime) and the change in area (proxy of cell volume) of single yeast cells subjected to 1 M NaCl. r = 0.06, p-value = 0.522. Δlifetime = (final lifetime - initial lifetime)/initial lifetime. Δarea = (final area - initial area)/initial area.

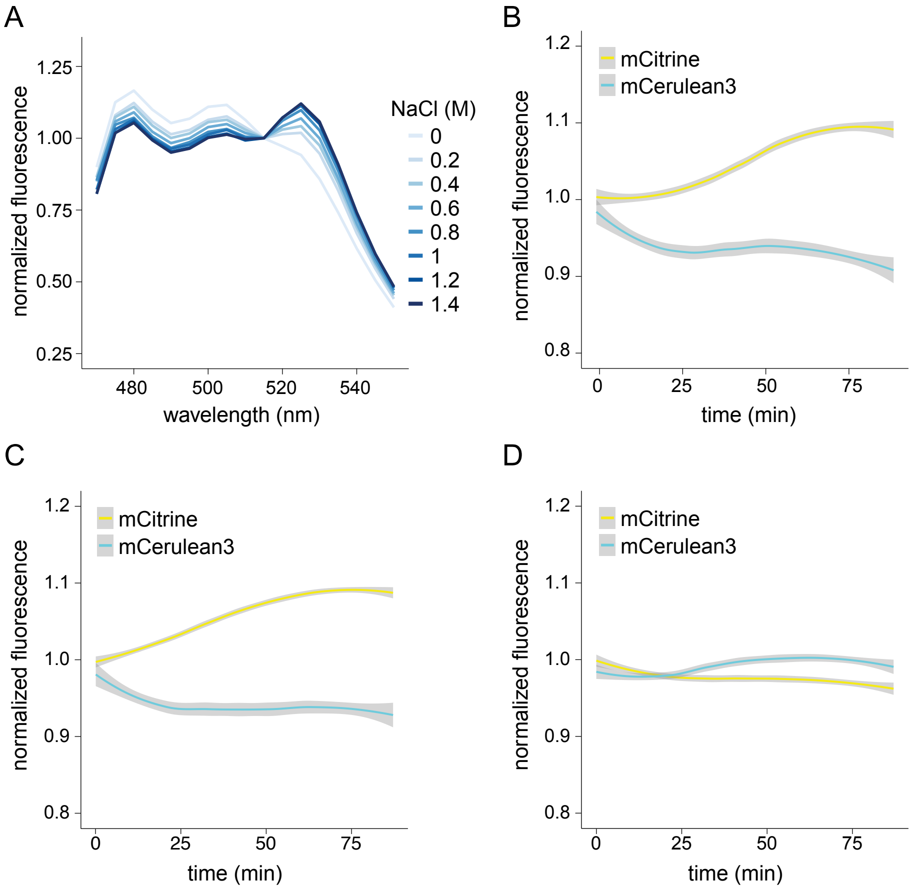


**Figure S8.** (A) Fluorescence emission spectra of NaCl-treated live *Escherichia coli* cells expressing SED1. Fluorescence values were normalized to the value at 515 nm. (B) Normalized fluorescence emission time course of the donor (mCerulean3) and acceptor (mCitrine) fluorophores in *Nicotiana benthamiana* leaf discs transiently expressing SED1, treated with 1 M sorbitol. n = 12 leaf discs. Mean ± SEM. One-way ANOVA. (C) Normalized fluorescence emission time course of the donor (mCerulean3) and acceptor (mCitrine) fluorophores in *Nicotiana benthamiana* leaf discs transiently expressing SED1, treated with 0.5 M NaCl. n = 12 leaf discs. Mean ± SEM. One-way ANOVA. (D) Normalized fluorescence emission time course of the donor (mCerulean3) and acceptor (mCitrine) fluorophores in *Nicotiana benthamiana* leaf discs transiently expressing SED1, treated with pure water. n = 12 leaf discs. Mean ± SEM. One-way ANOVA.

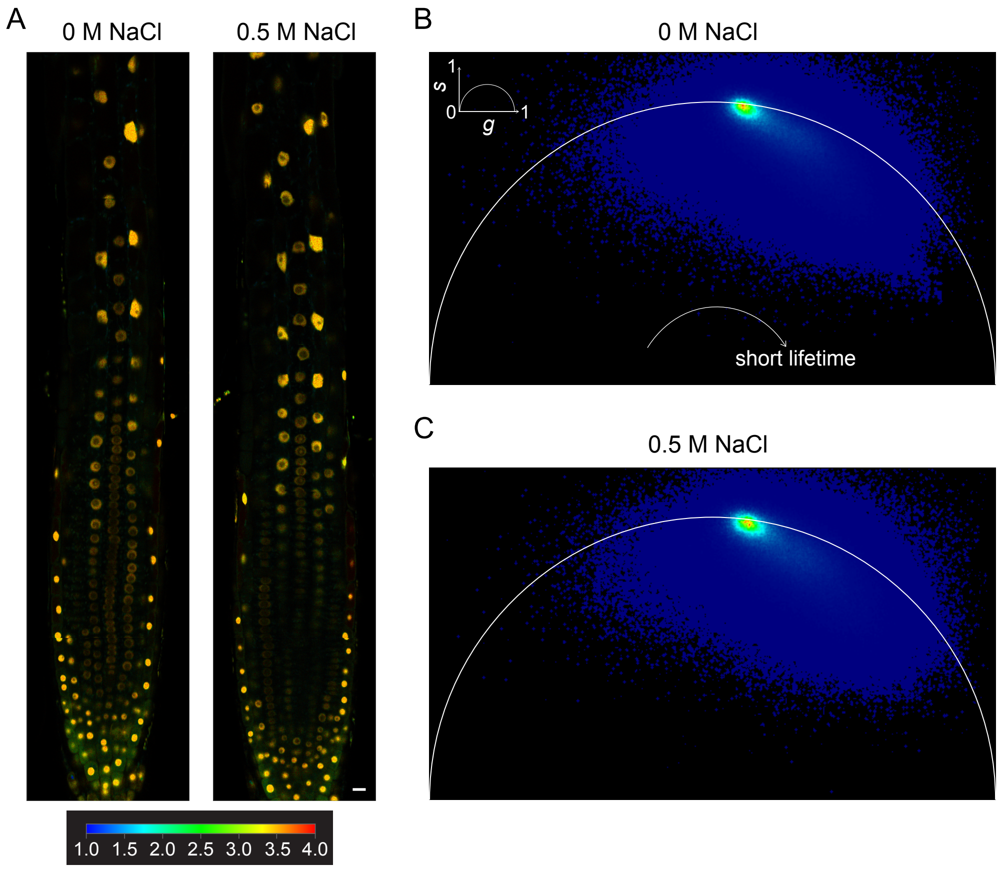


**Figure S9.** (A) Donor fluorescence lifetime images of *pUBQ10::nlsSED1* Arabidopsis roots imaged under 0 M NaCl or 0.5 M NaCl. Signal is localized to nuclei. Scale bar = 10 $\mu$m. Calibration bar represents the donor fluorescence lifetime in nanoseconds (ns). (B) Phasor plot of live *pUBQ10::nlsSED1* Arabidopsis root subjected to 0 M NaCl. (C) Phasor plot of live *pUBQ10::nlsSED1* Arabidopsis root subjected to 0.5 M NaCl. Signals shifted to the left side of the phasor plot represent longer fluorescence lifetimes, whereas signals shifted to the right side represent shorter fluorescence lifetimes.

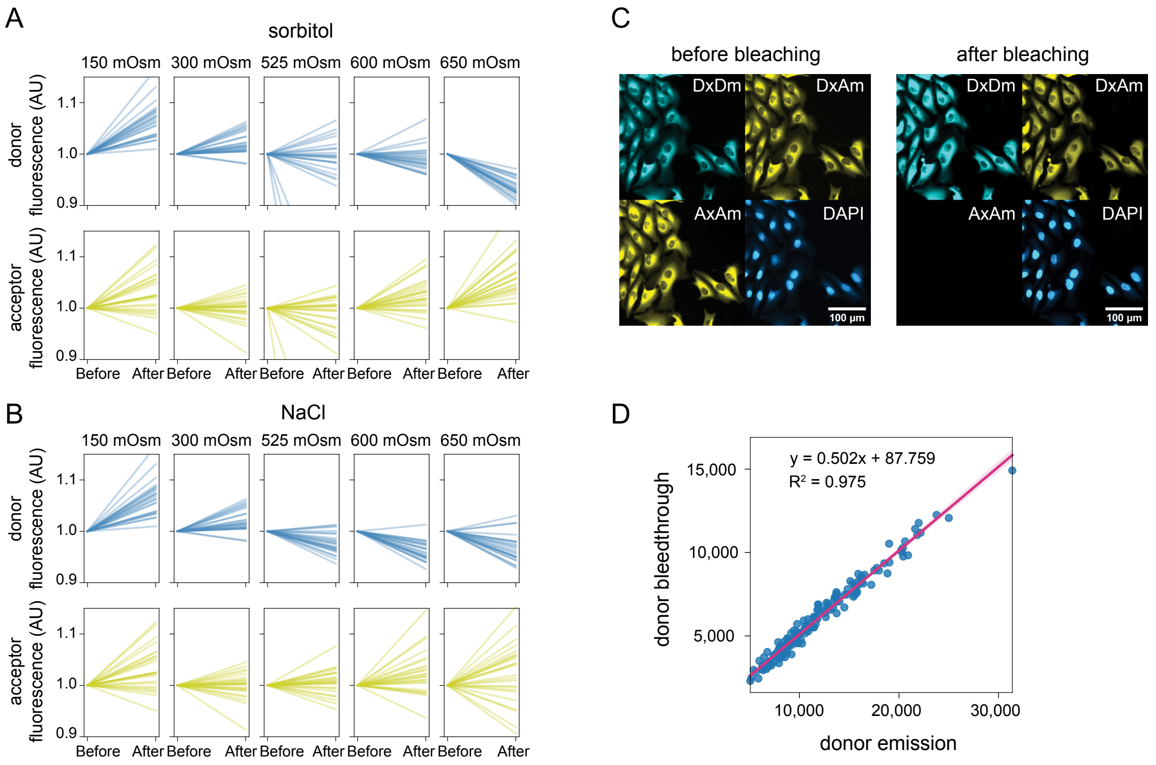


**Figure S10.** (A) Donor and acceptor fluorescence trajectories before and after the treatment of SED1-expressing U-2 OS single cells with sorbitol at the indicated osmolarities. (B) Donor and acceptor fluorescence trajectories before and after the treatment of SED1-expressing U-2 OS single cells with NaCl at the indicated osmolarities. (C) Image sample of live U-2 OS SED1-expressing cells imaged by donor (Dx, 430 nm), acceptor (Ax, 511 nm), or DAPI (385 nm) excitation. To correct for donor bleedthrough, cells were imaged (before bleaching), the acceptor was then photobleached, and the cells were imaged again (after bleaching). (D) A correlation plot of donor (DxDm) against acceptor (DxAm) emission was used to determine the bleedthrough correction. Multiple wells were imaged and measurements of all cells present in the plain of view were taken from the bleached images.

**Video S1.** Timelapse of donor fluorescence lifetime of single yeast cells expressing SED1 exposed to 1 M NaCl treatment at time 0 min. Scale bar = 10 $\mu$m.
